# Estimating Sparse Neuronal Signal from Hemodynamic Response: the Mixture Components Inference Approach

**DOI:** 10.1101/2019.12.19.876508

**Authors:** Anna Pidnebesna, Iveta Fajnerová, Jiří Horáček, Jaroslav Hlinka

## Abstract

The approximate knowledge of the hemodynamic response to neuronal activity is widely used in statistical testing of effects of external stimulation, but has also been applied to estimate the neuronal activity directly from functional magnetic resonance data without knowing the stimulus timing. To this end, sparse linear regression methods have been previously used, including the well-known LASSO and the Dantzig selector. These methods generate a parametric family of solutions with different sparsity, among which a choice is finally based using some information criteria. As an alternative we propose a novel approach that instead utilizes the whole family of sparse regression solutions. Their ensemble provides a first approximation of probability of activation at each timepoint, and together with the conditional neuronal activity distributions estimated with the theory of mixtures with varying concentrations, they serve as the inputs to a Bayes classifier ultimately deciding between the true and false activations.

As we show in extensive numerical simulations, the new method performs favourably in comparison with standard approaches in a range of realistic scenarios. This is mainly due to the avoidance of overfitting and underfitting that commonly plague the solutions based on sparse regression combined with model selection methods, including the corrected Akaike Information Criterion. This advantage is finally documented on fMRI task dataset.

## 1. Introduction

The most common approach to fMRI measurement is using the Blood Oxygenation Level Dependent (BOLD) contrast, see Ogawa et al. (1990). Soon after its discovery, it was shown that it reliably reflects changes in local brain activity related to spontaneous or stimulation-triggered cognitive processing, see Ogawa et al. (1992).

However, the BOLD signal is only indirectly related to the underlying neuronal activity. As its name suggests, it reflects the changes in the blood oxygenation level at a given location. These clearly appear only after the triggering neuronal activity, and constitute a considerably delayed and smoothed response to it that can be in first approximation modelled as the output of a linear filter represented by the so-called hemodynamic response function. Combined with a range of noise sources in fMRI measurements (Bianciardi et al., 2009), the detection of the appearance and timing of neuronal activation from the fMRI BOLD signal poses a serious signal analysis challenge.

The most common approach to deal with this is to impose strong prior information on the timing of the tentative neuronal activity, assuming that it amounts to a predefined time-course corresponding to the presence of particular sensory stimuli or other controlled experimental manipulation. Such a priori assumed activation time course is then convolved with a template hemodynamic response profile and used as a predictor in a linear regression model of the observed BOLD fMRI signal.

Yet there are still many scenarios when this approach is not suitable, typically when the precise timing of the presumed neuronal activity is not known as it is not perfectly locked to a predefined task paradigm, but the activations are highly dynamic, inter- and intra-individually variable, self-paced or ultimately spontaneously occurring. As rare as such experimental designs might occur in the field dominated by the use of regular block and event-related designs, having alternative tools available to analyse such less constrained fMRI BOLD data has previously been recognized as a valuable complement of the standard analysis methods Gaudes et al. (2013).

This alternative analytical approach is basically inverted with respect to the previously described – instead of convolving the expected neuronal activity with the hemodynamic response to obtain an approximate expected BOLD fMRI signal (and comparing this with the observed BOLD fMRI via linear regression), the observed signal is *deconvolved* to obtain an estimate of the underlying neuronal signal.

This ‘deconvolution’ task can be formulated as a linear regression problem that can be solved by various approaches; among those the so-called sparse regression approaches, that attempt to preclude overfitting of noise by inducing some penalties for marking too many activations in the data, have been mostly used. Some of the best known sparse regression methods are the LASSO (least absolute shrinkage and selection operator), see Tibshirani (1996), and the Dantzig Selector, proposed in Candes and Tao (2007). These methods output a parametric family of fitted models (parameterized by a free parameter determining the strength of the regularization and therefore the sparsity of the obtained solution). It is therefore important to use some suitable model selection procedure. The most known approaches are the Akaike Information Criteria (AIC) Akaike (1974) and Bayes Information Criteria (BIC) Schwarz (1978).

While these model selection criteria have been extensively and successfully applied in many fields, it is also known that the mentioned information criteria have their disadvantages in real-world applications. In particular, it has been shown that for a small sample size AIC tends to overfit the data, thereby giving too high a model order. On the other hand, the BIC-selected model may be underfitting the data, see e.g. Burnham and Anderson (2004). In such situations, the use of the adjusted Akaike Information Criterion (AICc) (Sugiura (1978), Hurvich and Tsai (1989)) was recommended to be used instead of AIC.

The application of sparse linear regression methods to the neuronal signal estimation problem is described in Gaudes et al. (2013), where the Dantzig Selector was used.

The authors compare AIC and BIC for the neuronal signal estimation task on the simulation study and show that AIC completely fails to estimate the sparse stimulus signal and overfits the fMRI time series. Therefore, in the aforementioned paper, the authors suggest using the BIC for model selection. However, the tested stimulus was very sparse in the simulation study, therefore we speculate that in the case of not-so-sparse activations, BIC might be underfitting the data.

In the presented work, we assess the performance of fMRI signal deconvolution methods including both the canonical LASSO as well as the Dantzig Selector regularization combined with the most popular selection criteria (AIC, BIC and AICc), arriving at a conclusion that their variable performance strength is far from perfect under some realistic scenarios, possibly due to nonrobustness of the model selection procedure. We therefore propose and present an alternative approach to the estimation construction based on the Bayes classifier and theory of mixtures with various concentrations, that appears to provide more accurate and robust results both in simulations and on a real data sample. The described algorithm is implemented in an updated version of software BRAD, whose basic functionality was described by Pidnebesna et al. (2018). The work is organized as follows. The practical motivation of the study and theory overview is given in Section 3. In Section 4 the new approach is proposed, methodology and simulation study are described. Finally, the real data application can be found in Section 6.

## 2. Data Model

We assume that we work with the measured BOLD signal *y* which can be modeled as a result of a hemodynamic response to the local brain activity *s*. In our model, the neuronal activity is represented by (a sparse) vector of activations; we assume there is no a priori information about stimulus timing. In line with the common fMRI BOLD modelling approaches, we assume that the measured signal is a result of the convolution of the neuronal activity and the hemodynamic response function (HRF), in mathematical notation:

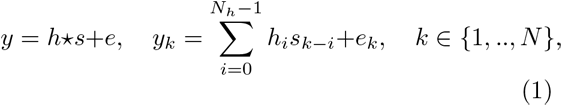

where *k* ∈ {1, .., *N*} indexes the time *\** denotes the convolution, *y, s, e* are *N* × 1 vectors denoting the measured signal, the underlying brain activity signal and the observational noise, respectively. The vector *h* denotes the HRF, where the length *N*_*h*_ is smaller than the length of the measured signal *N*_*h*_ < *N*. Here we assume that *e* is white noise, which is independent and identically distributed in each time point. In the matrix form, the model can be rewritten as

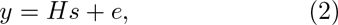

where *H* is the Toeplitz convolution matrix of size *N × N* corresponding to the HRF *h* (see Gray (2006))

In practice, the BOLD signal values output by the MRI scanner have a generally arbitrary scale and offset. Due to potential spontaneous neuronal activity fluctuations, establishing a proper baseline corresponding to ‘no substantial activity’ is a methodologically and even conceptually problematic task. Indeed, some level of ongoing ‘baseline’ activity can be expected, while we are aiming to detect the deviations from this baseline. As estimating the baseline value from the data itself also poses additional challenges, we have taken the heuristic approach of shifting the signal so that its relative range (minimum and maximum) around the zero value corresponds to that of the convolution kernel; alternatively just re-scaling the signal to have minimum 0 and maximum 1 gives comparable results. This is practically equivalent to assuming that the underlying (sparse) neuronal signal contains only *positive* activations (albeit of unknown amplitude). Note that such assumption is common in the field and often explicitly used in the model and optimization procedure definitions, see e.g. Hernandez-Garcia and Ulfarsson (2011); Bush and Cisler (2013). Of course, delineating the limits of both its conceptual and practical validity would require much deeper discussion of the nature of brain activity.

## 3. Motivation/Theoretical background

### 3.1. Sparse Regression Methods

In this paper, we chose to assess the two most prominent sparse regression methods - the LASSO (least absolute shrinkage and selection operator) and the Dantzig Selector. LASSO estimator was proposed in Tibshirani (1996), and it is a regression analysis method that performs variable selection. The aim (using notation congruent with the previous deconvolution problem introduction) is to estimate a sparse vector of linear regression coefficients *s* in linear regression:

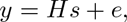

where *y* is the dependent variable, *H* is a matrix of regressors, and *e* is an additive i.i.d. Gaussian noise. Then the LASSO problem is formulated as

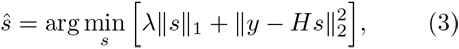

where *λ* ≥ 0 is a tuning parameter and ∥․∥_*p*_ is a norm in *L_p_*-space, i.e. 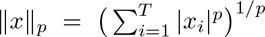.

Note that this is an instance of convex optimization and also of quadratic programming.

The Dantzig Selector was proposed in Candes and Tao (2007) as:

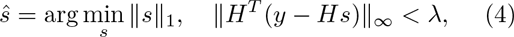

where *λ* ≥ 0 is a tuning parameter, ∥․∥_∞_ is a norm in *L*_∞_-space: ∥*x*∥_∞_ = sup_*i*_ |*x*_*i*_|. Similarity of LASSO and Dantzig Selector estimators were discussed, for instance, in James et al. (2009), Bickel et al. (2009), Asif and Romberg (2009).

### 3.2. Selection Criteria

In the model selection context, we assume that for given data, we obtain a set of candidate models. The selection criteria inform us about the relative quality of the candidate models, generally by finding a compromise between accuracy of the data fit and model parsimony. The model with the lowest value of the selection criteria among all of the tested models would be considered the best model.

The ***Akaike Information Criteria (AIC)*** is defined as

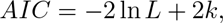

where *L* is the maximised value of the likelihood function of the model, and *k* is the number of estimated model parameters.

The ***Bayes Infromation Criteria (BIC)*** is defined as

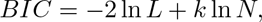

note that it also takes into account the sample size *N*.

In practice, the ***adjusted Akaike Information Criterion (AICc)*** is recommended to be used instead of AIC (see Hurvich and Tsai (1989), Burnham and Anderson (2004)) when *k* is large relative to sample size *N*. It is defined as

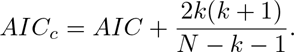

Clearly, that AICc converges to AIC for *N* → ∞.

Usage of these criteria for LASSO problems is discussed in Zou et al. (2007). For LASSO and Dantzig Selector, AIC and BIC have the form

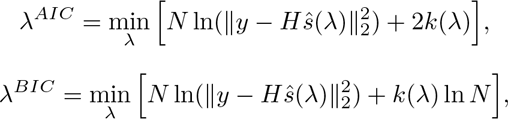

where *ŝ*(*λ*) is the solution of (3) and (4), that corresponds to *λ*, *y* is the measured signal, *H* is the matrix of regressors, *k*(*λ*) is the number of non-zero parameters in *ŝ*(*λ*) (see Zou et al. (2007)). Comparison of using the AIC and BIC to Dantzig Selector in deconvolution tasks in neuroimaging was discussed in Gaudes et al. (2013).

## 4. MCI

### 4.1. Basic idea: Bayes classifier

The main idea of our proposed approach is to utilize the information available in the observed data and the derived signals (namely the ordinal least square estimate of the brain activity and the family of regularized estimates) within a Bayesian inference tool that would separate the true activations from false positives due to observational noise.

The starting point is a basic estimate of the brain activity *ξ* = (*ξ*_1_, …, *ξ*_*N*_), *ξ*_*j*_ ∈ ℝ (think of e.g. the OLS solution *ŝ*_*OLS*_), that is typically not a sparse vector. We assume, that its (non-zero) elements correspond to activations caused by one of two possible reasons: neuronal response (true activation) or noise (false activation). Then we try to apply a naive Bayes classifier for two classes on the vector *ξ*.

Now, let us discuss the Bayes classifier in more detail. Let *g* : ℝ → {1, …, *M*} be the function (classifier), which returns the number of mixture component (class) *g*(*ξ*) ∈ {1, …, *M*} for each possible value of the observed characteristics *ξ* ∊ ℝ. In our case, there are two classes, i.e. *M* = 2, *g*(*ξ*_*j*_) ∊ {1, 2}, where *g*(*ξ*_*j*_) = 1 corresponds to the conclusion ‘*j*-th element falls to a true activation component’, and *g*(*ξ*_*j*_) = 2 corresponds to the conclusion ‘*j*-th element falls to a false detection component’. The Bayes classifier is of the form

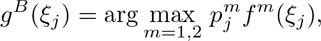

where *j* ∈ {1, …, *N*} indexes the classified elements of the vector, *ξ* ∊ ℝ are the basic (OLS) esti-mate values, 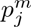 are the probabilities for *j*-th activation to fall to the *m*-th class, and *f*^*m*^ denotes the probability densities of the characteristic *ξ*, provided it falls within the class *m*. Thus, the classifier *g*^*B*^ (.) for each *j*-th activation compares the posterior probabilities of the element belonging to a neuronal activation or noise. Details and properties of such a classifier can be found, for instance, in Rish (2001) and Hand and Yu (2001).

However, true values of the probabilities *p*^*m*^ and the densities *f*^*m*^(*x*) are usually unknown a priori. Instead, we may need to obtain some estimates 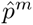 and 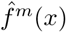 from data. We suggest two approaches to obtain such estimates. First, by using the Gaussian mixture model, described in McLachlan and Peel (2000). It is based on the assumption of the normal distribution of all mixture components. This approach is well-known, and its main advantage is a developed theory and existence of software tools for the automatic estimation of all the parameters of the model. For our purposes, it is important to estimate both of the needed elements, probability vectors 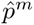 and component densities 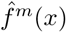.

The second approach is based on utilizing a model with varying concentrations (MVC) for estimating the component densities, and a heuristic procedure based on LASSO/Dantzig Selector algorithms for approximating the probabilities. Using this concept enables us, not only to disregard the assumption about the normality of the component distributions, but it also allows for utilizing element-specific prior probabilities – that are in the present scenario informed by the regularized regressions solutions. The detailed description of this concept is provided below and its block-scheme is shown in the left panel of Figure 1.

**Figure 1:**
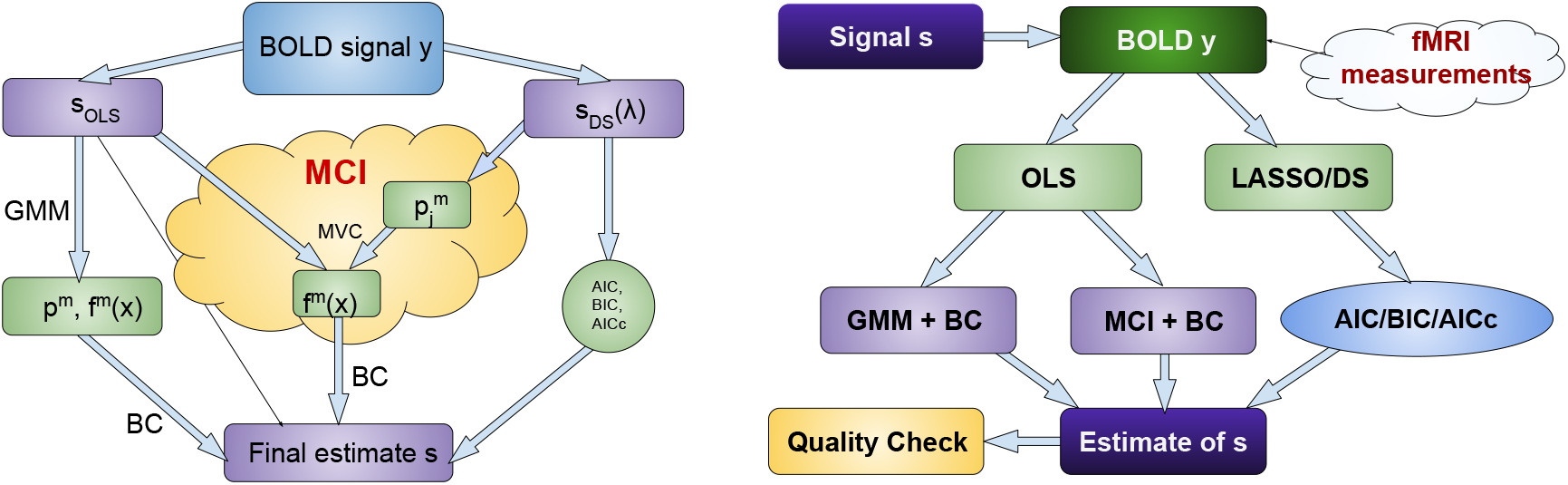
Left panel: Block-scheme of proposed MCI approach. Right panel: scheme of simulations.

### 4.2. MVC-based density estimates

In the classical finite mixture model, the mixing probabilities, and therefore the probability densities of all features *ξ*_*j*_, are supposed to be the same for all the observed objects *j* (McLachlan and Peel, 2000):

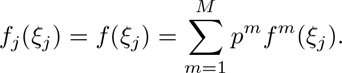

As a generalization of this approach, the model with varying concentrations (MVC) was proposed (Maiboroda and Sugakova, 2012). According to this model, mixing probabilities 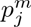, *m* ∈ {1, …, *M*}, *j* ∈ {1, …, *N*}, vary for different objects *j*, while still satisfying the condition: 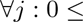 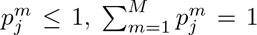. Then the distribution for the *j*-th object can be modelled as

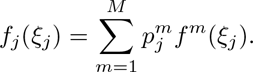

In MVC, the mixing probabilitiesFor both mentioned estimators, 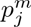 are sup-posed to be known, whereas the densities of components *f*^*m*^ are assumed to be unknown and estimated from the data. For estimation of the parameters, we use the kernel estimate

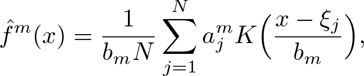

where *K*(.) is a kernel function and *b* is the kernel width and 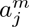 are the weights, aimed to distinguish the *m*-th component and suppress the influence of other components on it (see Maiboroda and Sugakova (2012) for details). The parameters are obtained using estimates developed in Maiboroda (2008) and Doronin and Maiboroda (2015). Classification of components of mixture are discussed in Sugakova (2006) and in Autin and Pouet (2012).

In particular, when one denotes the inner product of two vectors *a* = (*a*_1_, …, *a*_*N*_), and *b* = (*b*_1_, …, *b*_*N*_) by 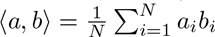 and defines the matrix

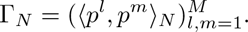

Note that this matrix is further assumed to be invertible, with *γ*_*km*_ being the *km*-minor of Γ_*N*_. Then the weights are given by:

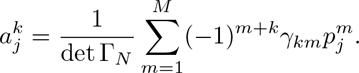

For the specific case of two-components, the weights 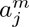 are given by:

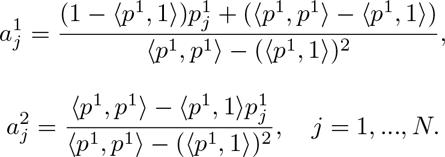

The kernel width *b*_*m*_ is estimated using the Silverman’s rule-of-thumb (see, for example, Hardle et al. (2006)):

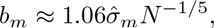

where 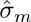 and 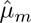 are the estimated standard deviation and mean of the *m*-th component given by 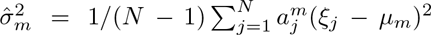, and 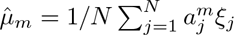, respectively.

### 4.3. Probability estimates

As it was described above, for the classification of signal values *s*_*j*_, we also need to set the probability 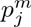 that it belongs to the true or false activations class. In this section, we propose a heuristic procedure based on the numerical solutions of the optimisation problems with a regularisation parameter. Particularly, we work with the homotopy algorithms for LASSO (3) and Dantzig Selector (4) estimators.

Let us denote *s*(*λ*) such solution of (3) (or (4)) that corresponds to a parameter’s value *λ*. Then, the support of *s*(*λ*) and its sign vector can be defined as

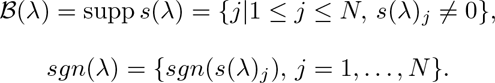

For both mentioned estimators, the particular solution *s*(*λ*), its support 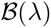 and sign vector *sgn*(*λ*) are specified by the parameter value *λ*. For the LASSO problem, it was shown in Efron et al.(2004), that for a given response vector *y*, there is a finite sequence of *λ*’s, called transition points (see Zou et al. (2007)),

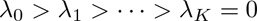

such that:

- for all *λ* > *λ*_0_, solution of the optimization problem (3) corresponds to the trivial solution, i.e. its support is the empty set 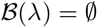;
- the value *λ*_*K*_ = 0 corresponds to the ordinary least squares solution;
- for *λ* values from the interval *I*_*m*_ := (*λ*_*m*+1_, *λ*_*m*_), the solutions of (3) are constant with respect to *λ* in terms of their supports and sign vectors;

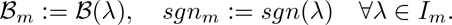

In other words, there is a sequence 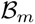, *m* = 0, …, *K* of subsets of indices *j, j* = 1, …, *N* that correspond to nonzero coordinates of *s*(*λ*) for ∀*λ* ∈ *I*_*m*_. Thus, for each *j* = 1, …, *N* there is a set of intervals *I*_*m*_, where the *j* lies in the corresponding solution support 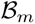:

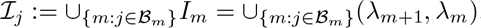

Further, let us denote the union of all intervals *I*_*m*_ as 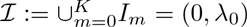. Then we propose to estimate the probability *ω*_*j*_ that *j*-th activation comes from the true neuronal signal as

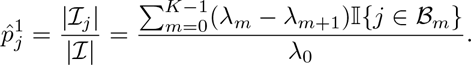

This approach corresponds to the idea, that more frequent appearance of non-zero activation estimate in the solution sets means higher probability for this moment to correspond to a true neuronal activation. While the probabilistic interpretation of this ‘relative occurence in the solution set’ is clearly heuristic, it is reasonably motivated as the LASSO operator is assumed to detect true variables among the noise.

Numerically, the set of transition points and corresponding sets of solutions for LASSO can be obtained by the homotopy algorithm LARS, presented in Efron et al. (2004). For Dantzig Selector, there is an available homotopy algorithm Primal-Dual Pursuit, which was presented in Asif and Romberg (2009). Note that the Primal-Dual Pursuit produces a finite sequence of so-called critical values of the parameters, that includes the points of changes in the support or sign vectors of the solutions; without loss of generality, these critical values can be considered as the transition points.

Thus, the probability 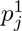 for *j*-th activation to come from true activation, can be approximated based on the numerical solution of the optimisation problem with regularisation parameters. Then, the probability 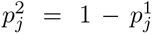, and the densities *f* ^1^(.), *f* ^2^(.) can be estimated. Finally, this information is enough to build the classifier and separate the components of vector *ξ* on true and noisy activations.

## 5. Simulation study

To assess the standard methods as well as the newly proposed methods, we use numerical simulations. We generate a signal *y* = *s* * *h* + *e* using HRF *h*, while parametrically varying the the properties of the signal *s* and noise *e*. Then, LASSO and Dantzig Selector with AIC/AICc/BIC selection criteria are used to estimate the input signal *s*. These estimates compared with two approaches based on Bayes Classifier: GMM and the proposed method MCI. To assess the quality of the obtained estimates, several measures are used. General scheme of simulations is shown in right panel of Figure 1.

The HRF kernel is obtained using the function *spm_hrf(.)* in the SPM toolbox (The Wellcome Dept. of Cognitive Neurology, University College London), setting scan repeat time (RT) to 2.5s. To define the level of added noise *e*, we use the Signal-to-Noise Ratio (SNR), 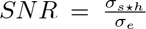, where *σ*_*s*\**h*_ and *σ*_*e*_ are standard deviation of BOLD signal and noise, respectively. For our simulations, we use several values of SNR in range [1.5, 4.5].

The input signal *s* was simulated as a vector of the length *N*, that consists of *K* non-zero activations (value 1) and *N − K* zero values. Positions of non-zero elements are chosen uniformly in the range [1, …, *N*].

The value of the selection criteria depends on the length of the signal *N* and number of non-zero elements *K*. Thus, we explore the estimates for several different noise levels and multiple numbers of true non-zero coefficients.

We denote the number of true negatives as *TN*, true positives as *TP*, false negative and false positive as *FN* and *FP*, respectively. To assess the model quality, we use several measures: Jaccard index, sensitivity, and specificity. The Jaccard index (Jaccard, 1901) is a similarity measure, defined as

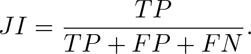

*JI* takes values in the range [0, 1], where 1 corresponds to perfect agreement. This similarity coefficient was also discussed in Gower (2004), Kosub (2016), or Willett et al. (1998).

Sensitivity, or true positive rate, is the probability of a positive test decision given that the true value is positive. Specificity is a probability that a test result will be negative when the condition is not present (true negative rate):

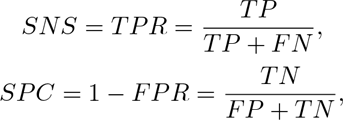

*SNS* and *SPC* take values in range [0, 1], where 1 corresponds to perfect specificity/sensitivity.

### 5.1. Numerical results

In this section, we present the numerical results of the simulation study. Figure 2 shows an example of a simulated signal and the obtained estimates. The red peaks are marks of the non-zero components of the simulated signal, the blue line is a noisy convolution of the corresponding simulated vector, thick black peaks are the obtained estimates (by the MCI DS method).

**Figure 2:**
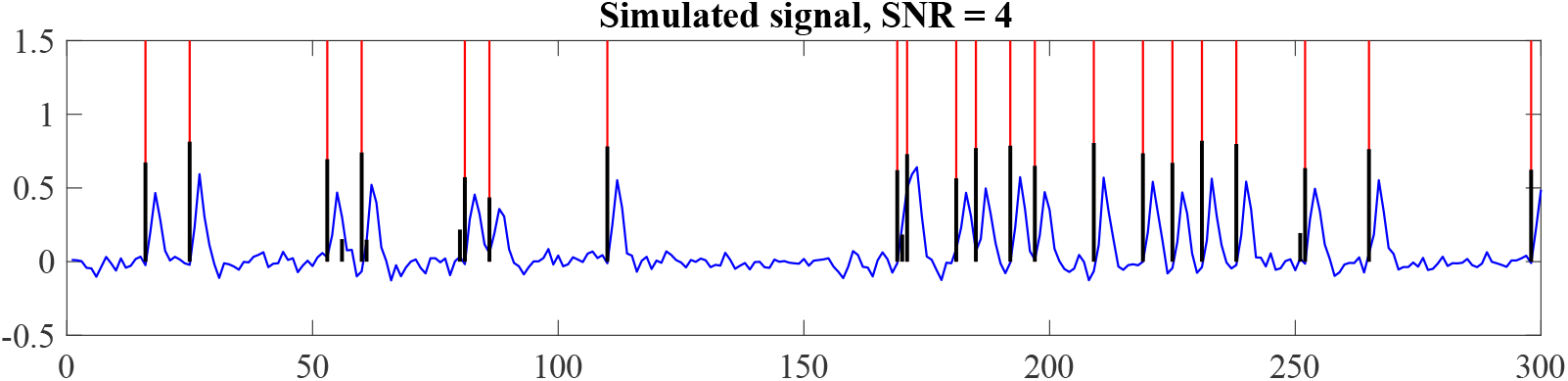
Example of a simulated and estimated signal. The simulated neuronal signal is shown in red, the simulated BOLD signal is shown in blue, the estimated neuronal signal is shown with thick black lines.

Our simulations show that DS/LASSO with AIC/BIC tends to mark almost all possible elements as non-zero, in other words, extremely over-estimates the signal. Figures that demonstrate corresponding results can be found in the Supplementary material. Thus, further results in this section are presented for DS AICc, MCI DS, and Bayes classifier based on GMM components (BC GMM).

Figure 3 shows the averaged results for 100 simulations for varying noise levels and activation density. The matrix contains 7 noise levels, increasing from top to bottom on the *y*-axis, and a different number of peaks on the *x*-axis (from the smallest to the highest values).

**Figure 3:**
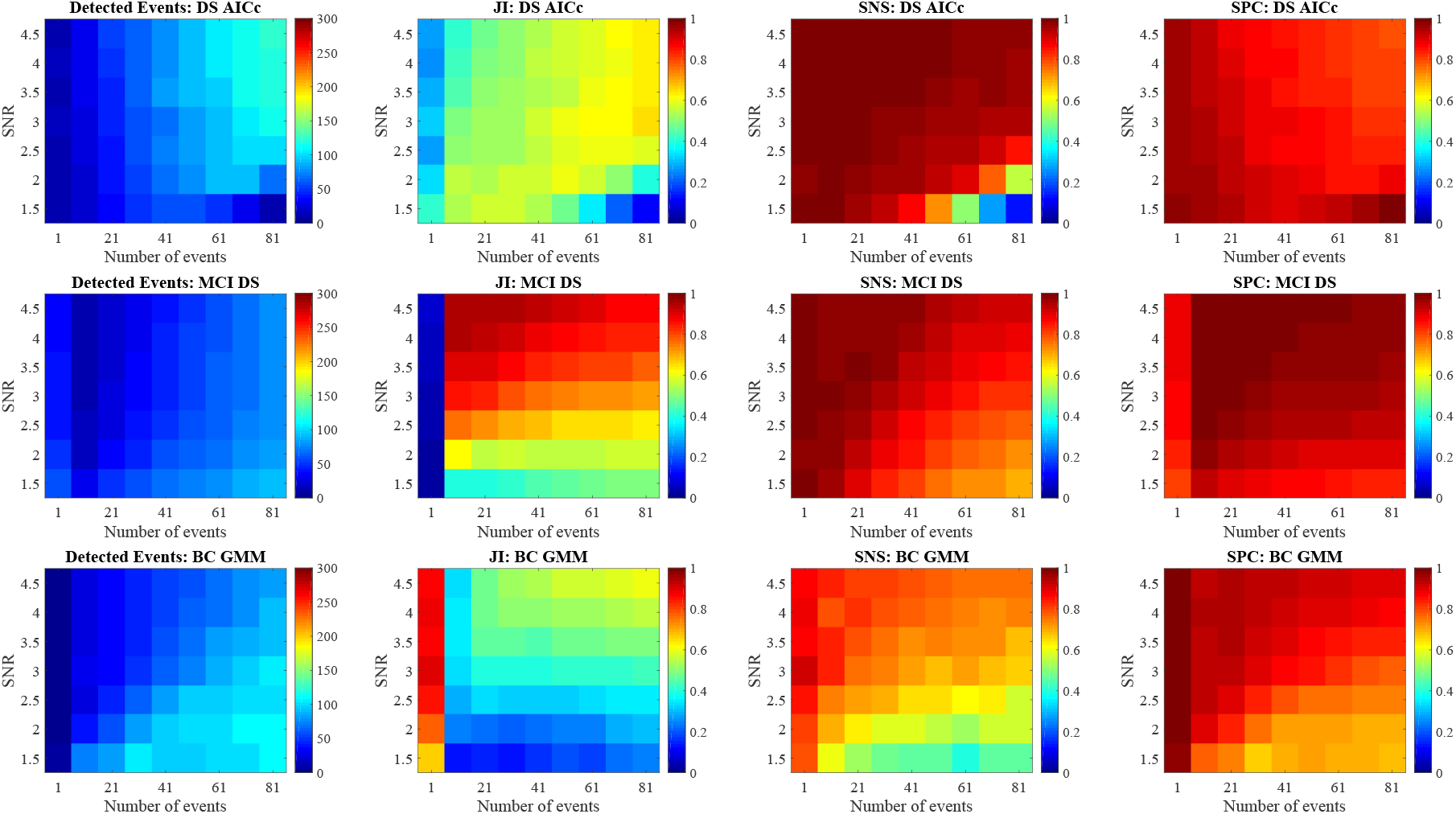
Comparison of the deconvolution methods on simulated data on the number of detected events, Jaccard index (JI), sensitivity (SNS), and specificity (SPC). Time series length is *N* = 300, the number of non-zero peaks is shown on the *x*-axis, the noise level on the *y*-axis. Estimation methods are shown in different rows: DS AICc, MCI DS and BC GMM.

Each row of the Figure corresponds to one of three compared methods: DS AICc, MCI DS and BC GMM. Every column shows one of the characteristics of the estimates: number of detected nonzero events, Jaccard index, sensitivity and specificity.

Clearly, BC GMM works comparably good only for very sparse signals. Also, this method tends to over-estimate the signal for higher noise.

The other two methods show generally better results. DS AICc tends to slightly over-estimate the number of detected events for higher SNR and less sparse signals. Otherwise, the method performs relatively well and gives stable results for the presented parameters. The Jaccard index is typically about 0.6, which is much better than the results of BIC or uncorrected AIC.

Concerning MCI DS, it fails for extremely sparse signal (only one activation from 300 points) for any SNR, which happens due to inability to estimate the distribution of true activations using only one point. However, for other parameters settings it shows good results. In particular, the quality of obtained estimates by MCI DS is even better than that of DS AICc, mainly in terms of the Jaccard index.

## 6. Real data example

### 6.1. Subjects

The real data example includes the data of 56 subjects (24 males and 32 females, age = 33.66 ± 10.97) that performed fMRI recording with n-back task as a part of a complex study with repeated psychiatric, neuroanatomical and neuropsychological evaluation. All subjects have been recruited for the study based on the following exclusion criteria: history of mental disorder, history of neurological disorder, presence of any artificial objects that would interfere with the magnetic resonance imaging. All volunteers signed Informal consent approved by the Ethic committee of National Institute of Mental Health (Czech republic). The data set of 89 MRI sessions in total was analyzed, including data from repeated sessions (6 months apart) in 25 subjects (2 visits for 17 subjects and 3 visits for 8 subjects).

### 6.2. N-back task

The n-back task is a continuous performance task commonly used for assessment of working memory performance (Gazzaniga et al., 2009). The subject is presented with a sequence of stimuli, and the task requires indicating when the current stimulus matches the one from n steps earlier in the sequence. If the task load factor n equals 2 or more, the working memory buffer needs to be updated continuously to keep track of what the current stimulus must be compared to. The spatial n-back task paradigm applied in this study presents the stimulus (blue square presented on black background) that appears in one of the 9 possible positions 3×3 matrix) on the screen during each turn. The tasks consisted of two alternating conditions: 0-back and 2-back. Each condition was presented in four sessions (eight sessions in total, sequence order: 0-2-0-2-0-2-0-2). The paradigm required the subject to recall the spatial position of the blue square a) in initial position for the 0-back session and b) two turns back for 2-back session, and press the button during the experiment based on these rules specific for each of the n-back conditions. Importantly, for the purpose of this study, we did not distinguish between the two conditions (0- and 2-back) or the correctness of the subjects’ responses, we simply considered/analysed all events of button presses recorded for each subject. For each subject, we have the button presses timing, which is assumed to correspond to the true activations of the motor cortex. As all subjects had the same experimental design, we can also carry out a group level analysis, setting as group ground-truth as button presses present in at least 80% of subjects (71 of 89)] and to average the measured BOLD signal in order to decrease noise. Also, the analysis of each subject separately was done. On average the subjects responded by button press 44.63 times (SD = 6.18, min = 27, max = 58). Most of the responses were correct, on average 5 of the responses were incorrect.

### 6.3. fMRI acquisition and preprocessing

Brain images were obtained using Siemens Prisma 3T MR machine. Functional T2*-weighted images with BOLD contrast were acquired with voxel size 3×3×3 mm, slice dimensions 64×64 voxels, 32 axial slices, repetition time 2000 ms, echo time 30 ms, flip angle 70°.

The data pre-processing was done using SPM8 toolbox (The Wellcome Dept. of Cognitive Neurology, University College London) in Matlab (The MathWorks, Inc.). In order to minimise head motion effects, functional volumes were spatially realigned, and the slice-time correction was used to fix acquisition delays. This was followed by normalization of functional volumes into the standard anatomical space using a template provided by SPM toolbox and spatial smoothing with 8 mm FWHM kernel.

An independent component analysis was performed in order to extract time series corresponding to the primary motor cortex, which was done using GIFT toolbox (MIALAB, Mind Research Network) in Matlab. The number of independent components was estimated using the minimum description length criteria. Independent components were decomposed with an Infomax algorithm (see Bell and Sejnowski (1995)) with default settings. A component representing the primary motor cortex was selected by visual inspection, see Figure 9. Its time series was further analysed. Moreover, on each of the mentioned time series the outlier correction was carried out using Tukey’s test (see Tukey (1977)) and the re-scaling discussed in Section 2 was applied. To check that good correspondence of the stimuli and the estimated neuronal activations was not caused by any motion artefact related to the motor activity during the button press, we did the deconvolution for all the ICA components with all the methods and compared the results with stimuli. A significant match of the estimate to the real stimuli was found only for two components that overlapped with motor cortex; the unilateral component, that showed higher agreement with the stimulus, was selected for subsequent analysis.

### 6.4. Results

In this section, we compare the standard approaches DS and LASSO with the selection criteria AIC, AICc, BIC, and our method MCI DS and MCI LASSO. To test that we use a suitable HRF kernel, we have first applied the forward model using the standard HRF function available in the SPM software, as well as its versions shifted by 1 TR backward and forward. The theoretical signal obtained using the forward shifted HRF was more correlated with the real signal than the one using the original HRF (see Figure 4), suggesting that our data were more in line with this shifted HRF. Note that the shape and delay of HRF varies in principle across subjects, brain regions as well as acquisition protocols, causing extra degrees of freedom in fMRI analysis, however estimating the HRF is outside the scope of the current paper, so we just fix it heuristically to the shifted version of the standard template.

**Figure 4:**
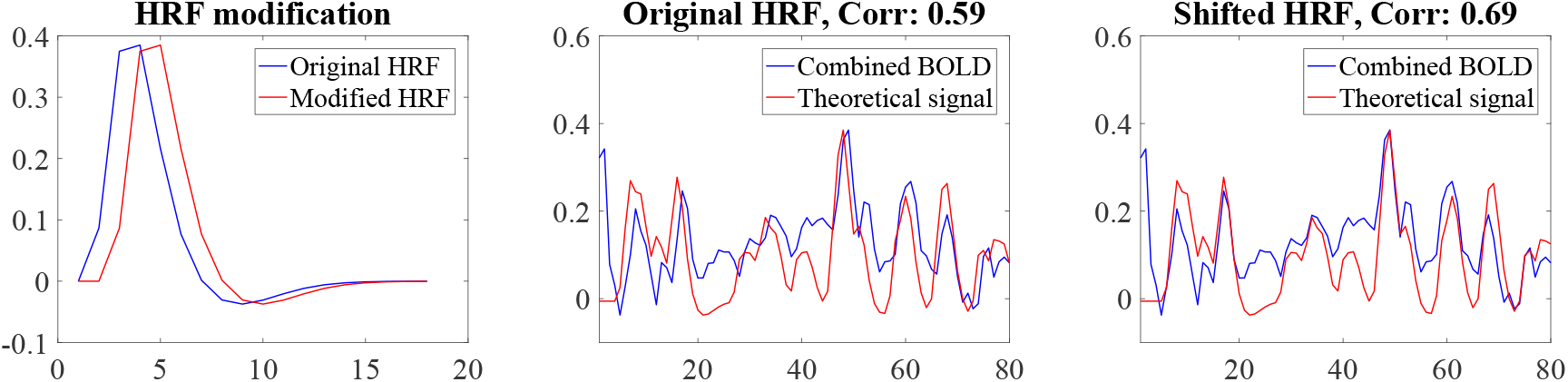
Selection of appropriate heamodynamic response function (HRF) using a forward model on real data from the N-back experiment. Left: the default and shifted HRF. Comparison of the averaged measured BOLD signal with the convolution of the true activations vector and the original HRF (middle) and shifted HRF (right).

In the next step, we applied all mentioned methods to compare the results using the quality measures discussed in Section 5. Then, Figure 5 shows the comparison of the random classifier with the discussed methods for original and shifted HRF. As can be seen, all of the methods for shifted HRF significantly differ from the random classifier, while for original HRF their results are comparable with it.

**Figure 5:**
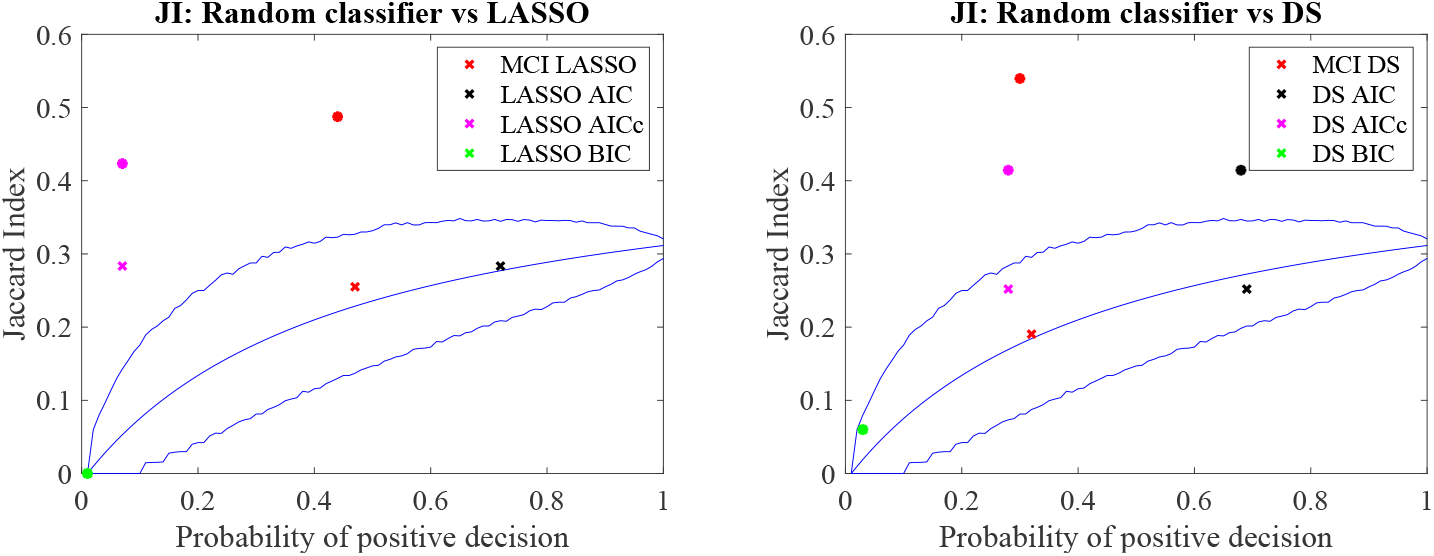
Comparison of the performance of the analysed methods with random classifier. For shifted HRF, the discussed methods show the results significantly different from the random one (confidence interval: [0.005,0.995]).

The results of activations signal estimation for the combined signal is shown on Figure 6. DS BIC-estimate is almost a trivial vector. It contains only true activations, but only two of them. Conversely, the DS AIC-estimate includes a lot of true detections, which gives the high sensitivity, but also it contains a lot of false detections. Remain two estimates both are better, and MCI DS shows higher values of all three quality characteristics.

**Figure 6:**
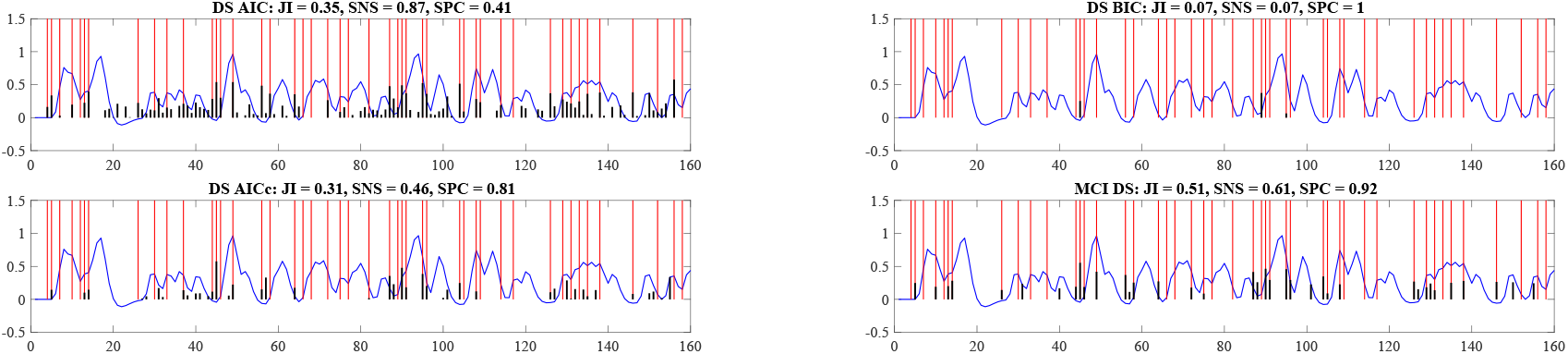
Application of the analysed methods to the group-level real data signal. The measured BOLD-signal is shown in blue, the theoretical activations in red, and the estimated activations in black.

Figure 7 shows the receiver operating characteristic curve (ROC-curve) for DS and LASSO approaches run on the combined signal. Numerical results for the shifted HRF are shown in Table 1. As it was described above, both of these methods generates the set of possible solutions on which the selection procedure should be applied. Thus, Figure 7 shows the ROC-curves with all discussed selection criteria. Also, the points corresponding to the MCI DS and MCI LASSO are marked on the graph, in order to compare the methods. The results demonstrate that the point corresponding to the MCI LASSO method lies close to optimal LASSO solution, and for DS the MCI performs much better than DS with the optimal selection technique.

**Table 1:** Comparison of the deconvolution techniques on the group-level real data signal with shifted HRF.

|  | JI | SNS | SPC |
| --- | --- | --- | --- |
| DS BIC | 0.07 | 0.07 | 1.00 |
| DS AIC | 0.35 | 0.87 | 0.41 |
| DS AICc | 0.31 | 0.46 | 0.81 |
| **MCI DS** | **0.51** | **0.61** | **0.92** |
| LASSO BIC | 0.00 | 0.00 | 1.00 |
| LASSO AIC | 0.42 | 0.94 | 0.45 |
| LASSO AICc | 0.11 | 0.12 | 0.96 |
| **MCI LASSO** | **0.49** | **0.78** | **0.73** |

**Figure 7:**
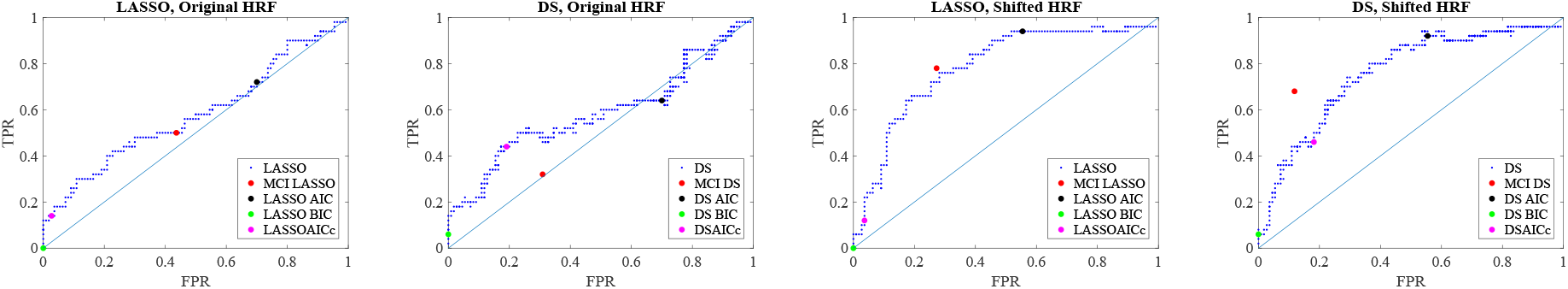
ROC curve of the Dantzig Selector and LASSO solutions for the group-level real data signal. Colored points specify the particular selection procedures.

Figure 8 shows the quality of estimates of each subject’s time series. It contains the marks of different colors for analysed methods. The black and green points that mark DS AIC and DS BIC methods form almost the same clouds and demonstrate that most of all estimates contain too many positive detections. The blue crosses mark the DS AICc and show a lot of estimates with too few true positive detections. The rest of the blue crosses forms a cloud that has an intersection with all other methods. The red color is used for MCI DS estimates, and this cloud lies apart from the DS AIC and BIC and demonstrates more appropriate estimates in terms of true and false positive rates. Numerical results are shown in Table 2.

**Table 2:**
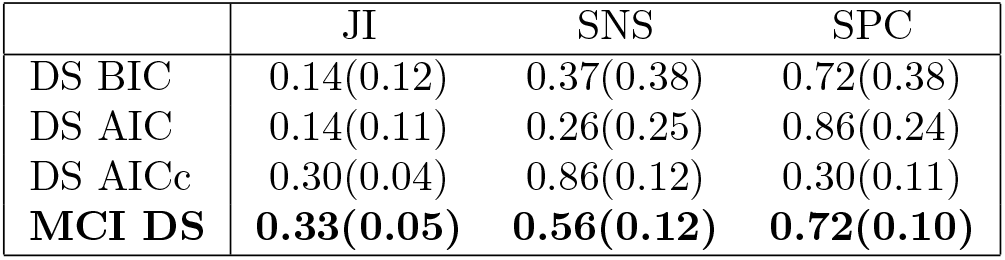
Summary performance characteristics (Jaccard index, sensitivity, specificity) for subject-level analysis. Mean (std) is shown for each method.

|  | JI | SNS | SPC |
| --- | --- | --- | --- |
| DS BIC | 0.14(0.12) | 0.37(0.38) | 0.72(0.38) |
| DS AIC | 0.14(0.11) | 0.26(0.25) | 0.86(0.24) |
| DS AICc | 0.30(0.04) | 0.86(0.12) | 0.30(0.11) |
| **MCI DS** | **0.33(0.05)** | **0.56(0.12)** | **0.72(0.10)** |

**Figure 8:**
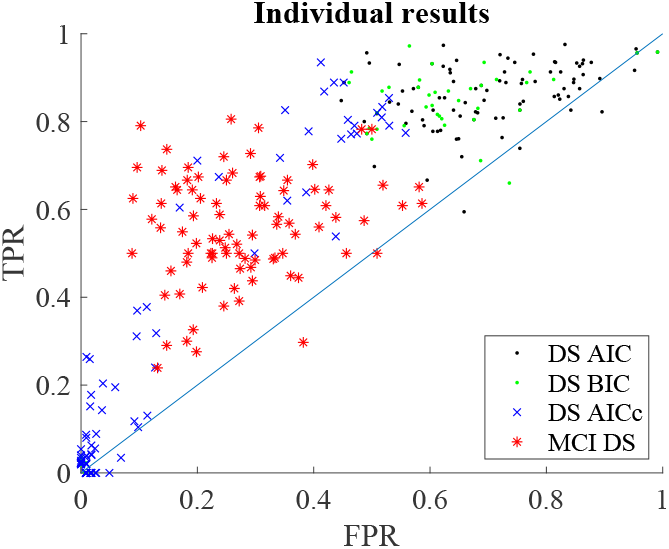
Comparison of deconvolution methods on individual-subject level.

## 7. Discussion and conclusions

We have introduced a method for improvement of the estimates of neuronal activity from BOLD signal. It works by applying an additional post-processing on the result of standard regularization-based deconvolution methods, such as LASSO and Dantzig Selector. Notably, it outperforms both the standard application of the Bayes information criterion and Akaike’s information criterion, as well as its improved variant.

The advantages of this method were demonstrated both in numerical simulations as well as in experimental data, where it provided more accurate estimates of timing of motor responses occurring during a complex cognitive task. This provides a proof of principle that the method would further improve the detection of neuronal activation in relation to processes that can not be directly recorded or controlled.

The main aim of the approach is to overcome the challenges of regularization methods combined with standard model selection criteria by combining the information contained in the solutions rather than select single one; this allows in particular to avoid the common overfitting or underfitting of the data and shows a good performance both in simulations and in a real dataset example. The future work would involve attempting its efficient combination with some other directions recently taken up in the field, namely spatial regularization, change sparsity, HRF estimation and further processing.

In more detail, the possible extensions and integrations include; the joint estimation of the HRF and neuronal activity, such as in the fused LASSO and Tikhonov regularization of the HRF Aggarwal et al. (2015), spatial structure modelling including regularization as in the work of Karahanolu et al. (2013), which was further extended in Farouj et al. (2017) to remove the need for a priori brain parcelation, or alternative penalty schemes in regularization such as combining total variation and negativity penalties as in Hernandez-Garcia and Ulfarsson (2011). Of interest could be the extension to application of such deconvolution using more complex data models, such as the bilinear model including neuronal activity autocorrelation, stimulation and modulation terms Penny et al. (2005); Makni et al. (2008), potentially in the context of network analysis.

As the outlined approach relies on being combined with an initial method providing a parameterized family of solutions (in this paper, we used the LASSO and Danzig classifier to provide examples of such families), we leave open the question of how it relates and could synergize with other families of approaches such as using the cubature Kalman filtering Havlicek et al. (2010), activelets (wavelets designed using prior knowledge on the heamodynamic response function form Khalidov et al. (2011)), spatiotemporal HRF deconvolution Aquino et al. (2014) or nonparametric hemodynamic deconvolution of fMRI using homomorphic filtering Sreenivasan et al. (2015). We can generally speculate that the key idea of combining model outputs using e.g. Bayesian approaches instead of crisp model selection within a constrained parametric family may be applicable to other schemes beyond classical regularization schemes such as LASSO, as long as these involve selection among model parameterized model families.

The obtained information on local neural activations can be of interest on its own such as in the application to study cortical response to unpredictable mental events as outlined in Gaudes et al. (2011), or possibly used for obtaining metaanalytic interpretation of the brain activation patterns Tan et al. (2017). Importantly, it has been proposed that functional connectivity in MRI is driven by spontaneous BOLD events detectable by deconvolution methods Allan et al. (2015); and the use of deconvolution as an intermediate step to estimate effective connectivity has been advocated Wu et al. (2013). Similarly, the dynamics of deconvolution-based brain activation patterns can be further explicitly modelled to fit sparse coupled hidden Markov models Bolton et al. (2018) characterizing the large-scale brain dynamics.

Last but not least, the development of deconvolution approaches is relevant not only for processing of most common fMRI BOLD signal but also in the context of functional near-infrared spectroscopy Santosa et al. (2019); Seghouane and Ferrari (2019) and quantitative susceptibility mapping Costagli et al. (2019). In general, this field demonstrates the fruitful interaction between experimental and technical challenges specific to each measurement modality or even particular type of experiment and the development of novel algorithmic and mathematical approaches, possibly applicable across disciplines.

## 8. Acknowledgement

We thank David Tomeček for preparing example fMRI data and Jessica Barilone for helping with language and style editing. This work was supported by Czech Health Research Council Project No. NV17-28427A and project Nr. LO1611 with a financial support from the MEYS under the NPU I program.

## 9. Supplementary material

**Figure 9:**
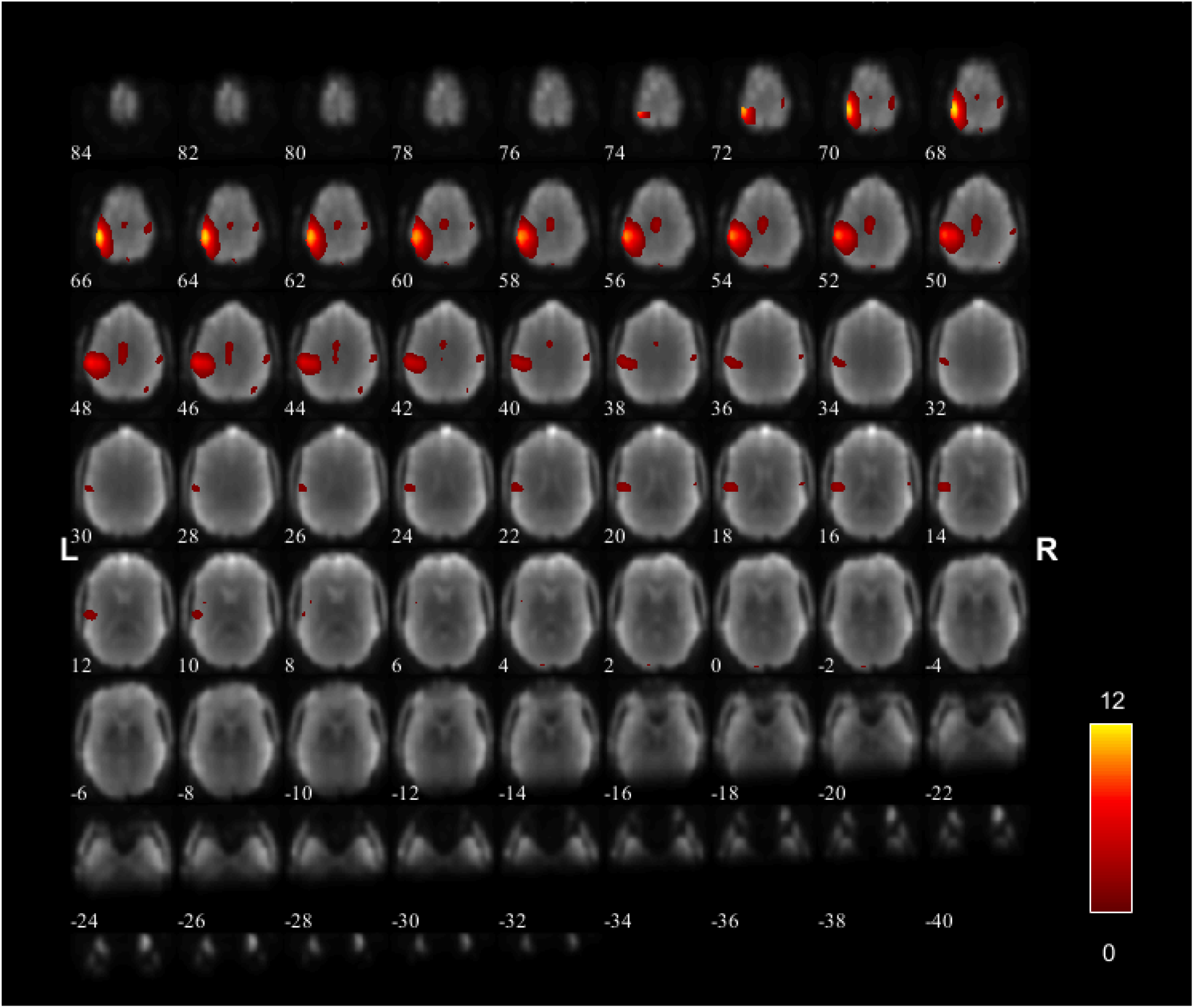
Primary motor cortex component.

**Figure 10:**
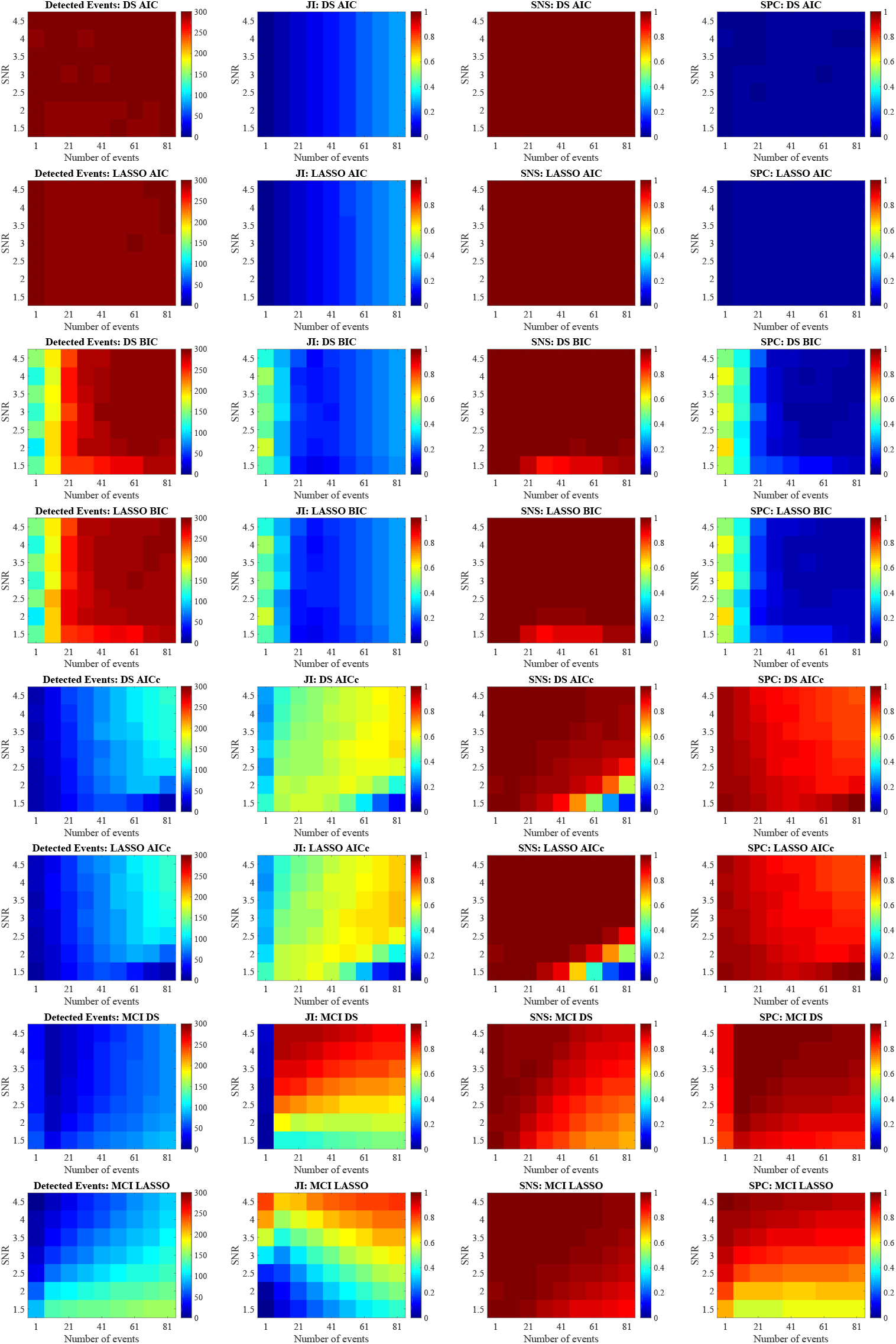
Comparison of DS and LASSO with standard selection criteria and MCI results.

